# Identifying and quantifying isoforms from accurate full-length transcriptome sequencing reads with Mandalorion

**DOI:** 10.1101/2022.06.29.498139

**Authors:** Roger Volden, Kayla Schimke, Ashley Byrne, Danilo Dubocanin, Matthew Adams, Christopher Vollmers

**Author notes:** Pacific Biosciences, Menlo Park, California 94025, USA. Genentech, San Francisco, California 94080, USA.

## Abstract

The Mandalorion tool, which we have continuously developed over the last 5 years, identifies and quantifies high-confidence isoforms from accurate full-length transcriptome sequencing reads produced by methods like PacBio Iso-Seq and ONT-based R2C2. In this manuscript, we introduce and benchmark Mandalorion v4 which further improves upon the already strong performance of Mandalorion v3.6 used in the LRGASP consortium challenge. By processing real and simulated accurate full-length transcriptome sequencing data sets, we show three main features of Mandalorion: First, Mandalorion-based isoform identification has very high Precision and maintains high Recall even when used in the absence of any genome annotation. Second, isoform read counts as quantified by Mandalorion show high correlation with simulated read counts. Third, isoforms identified by Mandalorion closely reflect the full-length transcriptome sequencing data sets they are based on.

## Introduction

In any eukaryotic cell, alternative splice site, transcription start site, and polyA site usage shape transcriptomes by enabling the expression of multiple unique isoforms for any one gene [1]. Understanding which isoform is expressed for a gene in a specific sample, be it a single cell or a bulk tissue, is crucial to understanding the biological state of that sample. Consequently, developing tools that gather a complete isoform-level understanding of the transcriptome within a sample is one of the main remaining challenges in genomics. With huge consortium projects such as the Earth Biogenome Project [2] now working on expanding our understanding of genomic diversity across species by sequencing and assembling new genomes, it is particularly important to develop isoform identification tools that do not depend on previously generated and curated genome annotations.

Tools designed to process data generated by the ubiquitous RNA-seq assay fail at the isoform-level analysis of transcriptomes [3–5] because RNA-seq uses short sequencing reads to sequence fragmented transcripts. Inferring or assembling accurate transcript isoforms from fragmented transcript sequences has proven to be extremely challenging [6]. However, while short-read platforms like Illumina are limited to read lengths of a few hundred nucleotides - much shorter than average transcripts, Pacific Biosciences (PacBio) and Oxford Nanopore Technologies (ONT) long-read platforms now routinely generate read lengths of tens of thousands of nucleotides - far longer than average transcripts.

Long-read platforms therefore made it possible to sequence transcripts end-to-end which in turn gave rise to the growing field of full-length transcriptome sequencing assays. To overcome the high error rate inherent to both PacBio and ONT platforms, the PacBio Iso-Seq method and the ONT-based R2C2 method generate consensus sequences from very long but error-prone raw reads. The millions of highly accurate end-to-end transcript molecule sequences that PacBio Iso-Seq [7,8] and ONT-based R2C2 [9–11] are producing have already been applied to bulk and single-cell transcriptome analysis. These studies have shown that data sets produced by these methods simplify the task of identifying isoforms dramatically, because if single accurate sequencing reads cover entire transcripts, inference or assembly of isoforms as done for fragmented data is unnecessary. Instead, isoforms can simply be defined by grouping and summarizing read alignments based on their features (i.e. alignment starts/ends and splice junctions).

Indeed, new computational tools have been developed or existing tools adapted to take full advantage of this new data type. However many of these tools rely heavily on previously generated genome annotations. As part of the LRGASP consortium, several isoform identification tools, including StringTie [3,12], IsoQuant [13], IsoTools [14], Bambu [15], FLAIR [16], FLAMES [17], TALON [18], and our own Mandalorion were compared by a group of independent evaluators to assess performance [19]. Mandalorion performed very well compared to these other tools, likely driven by high Recall when identifying non-annotated isoforms. Mandalorion also showed high Recall and Precision for the identification of SIRV spike-ins in R2C2 and PacBio data.

Here we introduce and benchmark version 4 of Mandalorion. Mandalorion v4 identifies isoforms with very high Recall and Precision when applied to either spike-in or simulated data with known ground-truth isoforms. Mandalorion v4 outperforms Mandalorion v3.6 (used in LRGASP) and StringTie v2.2.1 when identifying and quantifying isoforms. Further, Mandalorion v4 had a distinct performance lead over StringTie v2.2.1 when both tools were run entirely without annotation files. Running tools entirely without an annotation allowed us to evaluate performance within poorly annotated genomes like those of non-model organisms. We also show that Mandalorion not only accurately identifies isoforms but accurately quantifies isoform levels. Finally, by analyzing public PacBio Iso-Seq data, we show that isoforms identified by Mandalorion closely reflect the data set they are based on. Together, this establishes Mandalorion as an excellent choice when analyzing any high-accuracy long-read transcriptome data set.

## Results

### Mandalorion workflow

Mandalorion v4 accepts an arbitrary number of fasta/q files containing accurate full-length transcriptome sequencing data. While Mandalorion is generally platform and method agnostic, it has only been tested on high-accuracy end-to-end transcript molecule cDNA sequences which can be generated by either the ONT-based R2C2 method or the PacBio Iso-Seq method. Mandalorion is organized into several modules (A, P, D, F, and Q) which by default are all run sequentially (Fig. 1). The “A” module aligns reads using minimap2 [20]. Next, the “P” module converts and cleans (removing small indels) the resulting read alignments and the “D” module then processes the resulting clean read alignments locus by locus to 1) identify high-confidence splice sites, 2) group reads into junction-chains based on the splice sites they contain, 3) identify high-confidence TSS and polyA sites for each junction-chain, 4) define (potentially several) isoforms for each junction-chain based on TSS and polyA usage, and 5) generate consensus sequences for each isoform using pyabpoa [21]. The “F” module then aligns and filters the isoform consensus sequences (fasta) to generate isoform models (gtf and psl). Finally, the “Q” module quantifies the filtered isoforms across all provided input fasta/q files.

**Fig. 1:**
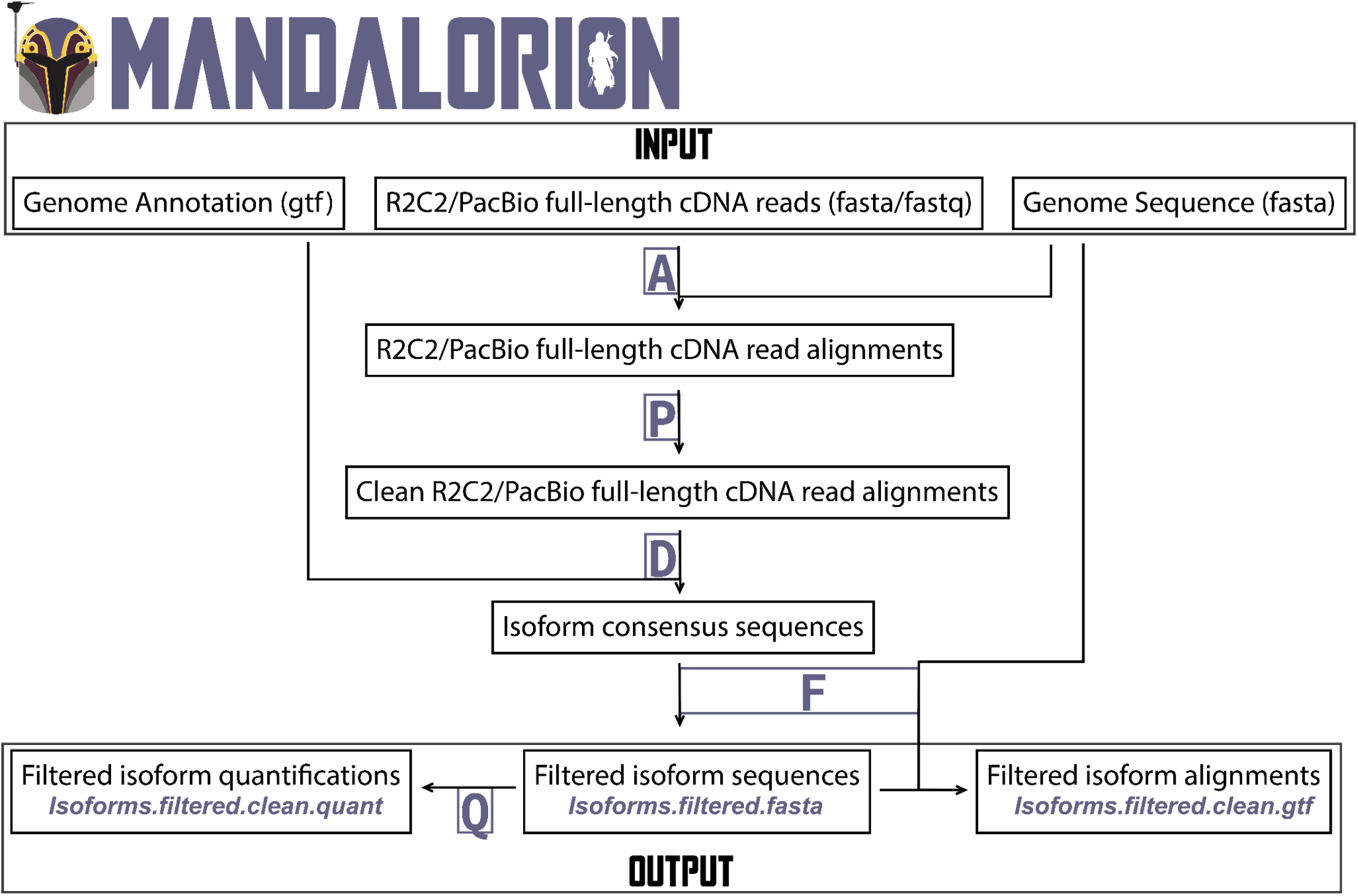
Mandalorion workflow. Input files, processing steps and output files are shown in a workflow diagram. Using several modules (A, P, D, F, and Q), Mandalorion aligns reads to a genome sequence (using minimap2), groups reads into isoforms based on those alignments, and generates a consensus sequence for each isoform (using pyabpoa). It then aligns these isoform sequences (using minimap2) and filters the isoforms based on those alignments.

### Evaluation of Mandalorion v4

We compared Mandalorion v4 and StringTie v.2.21 for the identification of isoforms from both simulated PacBio reads as well as real R2C2 and PacBio reads - whether or not a genome annotation was provided. We focused our comparison on StringTie v2.2.1 because it is heavily used and also does not require a genome annotation file as input which meant we didn’t have to use it in a way that wasn’t intended by the developers.

First, we ran Mandalorion v4 and StringTie v2.2.1 on mouse data produced or simulated by our group (R2C2 reads) and others (simulated and real PacBio Iso-Seq reads) for the LRGASP consortium and available from ENCODE. Second, we ran Mandalorion v4 and StringTie v2.2.1 on publicly available Universal Human Reference (UHR) PacBio Iso-Seq data. We also included Mandalorion v3.6 analysis but only for the analysis of simulated data because of its prohibitively long run time when no genome annotation is provided.

To compare the isoforms identified by Mandalorion v4 and StringTie v2.2.1 when run on each data set and in each condition, we used SQANTI [22] categorization. For simulated data, the entire known set of simulated isoforms served as ground truth, while for the real R2C2 and Iso-Seq LRGASP data, the set of known SIRV spike-in transcripts served as ground truth. For ground truth based analysis, isoforms scored as full_splice-match (FSM) to a ground truth isoform were considered as True Positives (TP) which in turn allowed us to calculate Recall (TP/(TP+FN)) and Precision (TP/(TP+FP)). For the analysis of the UHR data, which lacks a ground truth, we evaluated how isoforms identified by each tool and condition compared to each other and whether they reflected the actual read alignments.

#### Evaluating Mandalorion v4 isoform identification performance with annotation

With an annotation file available, Mandalorion v4 outperformed StringTie v2.2.1 when analyzing simulated PacBio Iso-Seq data. Mandalorion reached Recall and Precision of 87.96% and 92.23% compared to 77.91% and 91.86% reached by StringTie v2.2.1 (Table S1, Fig. 2A). Mandalorion v4 also represents a slight improvement over Mandalorion v3.6 used in the LRGASP challenge which reached Recall and Precision of 85.12% and 91.80%.

**Fig. 2:**
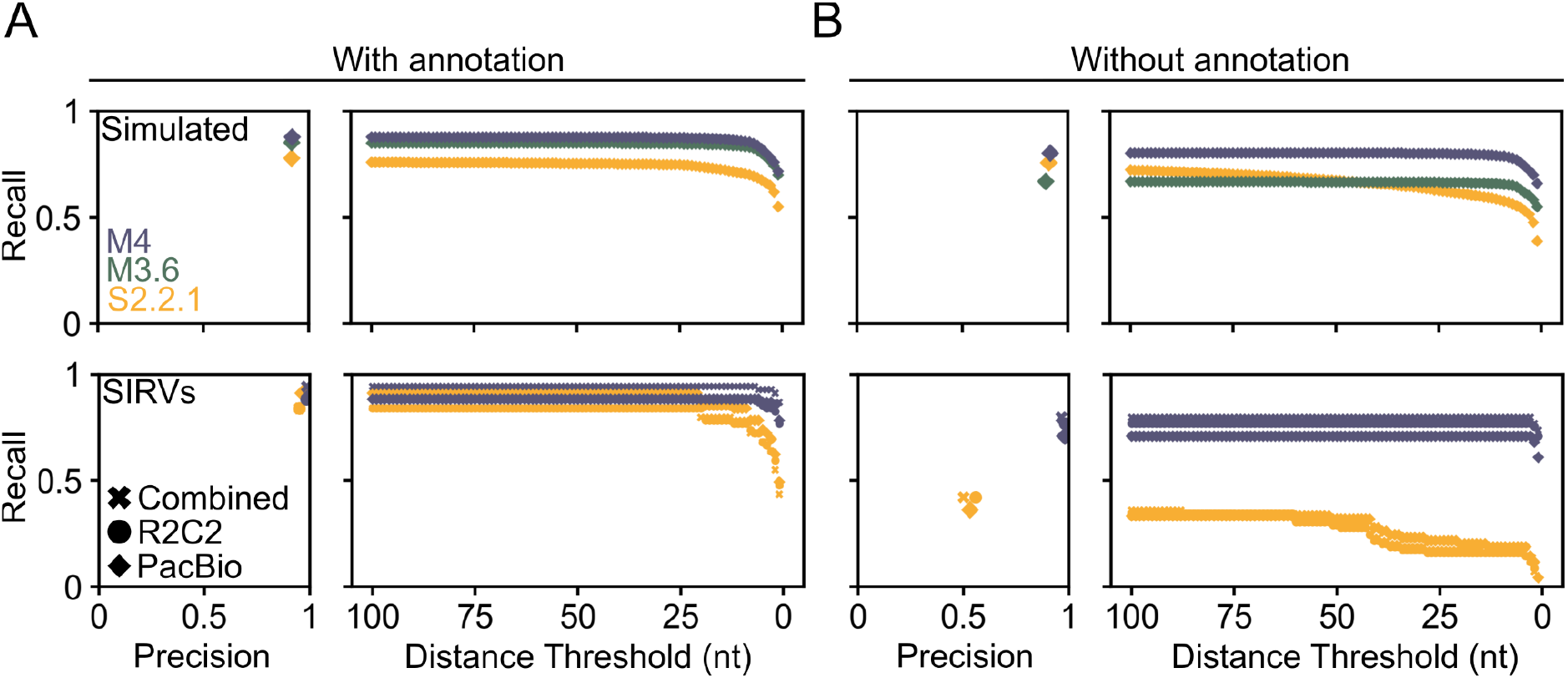
Mandalorion v4 outperforms StringTie v2.2.1 for isoform identification. Recall and Precision of Mandalorion v3.6 (M3.6), Mandalorion v4 (M4), and StringTie v2.2.1 (S2.2.1) are shown for Simulated PacBio data and real R2C2, PacBio, and combined R2C2 and PacBio (Combined) data. Recall and Precision are shown for tools run with (A) or without (B) annotation. Recall is also shown with distance thresholds applied.

Mandalorion v4 reached Recall and Precision of 89.86% and 100% when analyzing SIRV spike-ins in real PacBio data. Here StringTie v2.2.1 matched Mandalorion v4 performance very closely with Recall and Precision of 91.30% and 96.92% (Fig. 2A). For both tools, Recall and Precision for SIRV spike-ins in R2C2 and combined R2C2/PacBio data were very similar (Table S1).

For this initial analysis all full_splice-matches (FSM) to a ground truth isoform were considered as True Positives (TP). Next, we applied a distance threshold so that the ends of an identified isoform also have to fall within a certain distance of the ground truth isoform to be considered a True Positive. With increasingly stringent distance thresholds, the Recall of StringTie v2.2.1, declined more sharply than the Recall of Mandalorion v4 (Fig. 2A right).

This shows that, overall, when an annotation file was available, Mandalorion v4 outperformed or closely matched StringTie v.2.21 in both Recall and Precision for the identification of isoforms from both simulated PacBio reads as well as SIRV spike-ins in real R2C2 and PacBio reads.

#### Evaluating Mandalorion v4 isoform identification performance without annotation

Without an annotation file available, Mandalorion v4 reached Recall and Precision of 80.34% and 91.59% for simulated PacBio data compared to 75.64% and 90.86% reached by StringTie v2.2.1. In contrast to Mandalorion v4 (Recall 71.01%; Precision 98%), StringTie v2.2.1 reached much lower Recall and Precision without annotation (36.23% and 53.19%) for SIRV spike-ins in real PacBio data (Fig 2B).

To illustrate the performance difference between Mandalorion v4 and StringTie v2.2.1 on the complex loci the SIRV spike-ins are designed to represent, we visualized several SIRV loci. These genome browser style visualizations suggest that without genome annotation, StringTie combines the components of multiple separate transcript isoforms into False Positive isoforms (Fig. 3). In contrast, the only False Positive isoform identified by Mandalorion v4 was associated with spike-in transcript isoform SIRV503. Interestingly, while the isoform model generated by Mandalorion for this isoform lacked a 7 nucleotide terminal exon, the read-derived consensus sequence generated by Mandalorion for this isoform contained these 7 nucleotides. This highlights that even if a perfect consensus sequence is generated for an isoform, identifying very short terminal exons will continue to represent a formidable challenge for any long-read aligner.

**Fig. 3:**
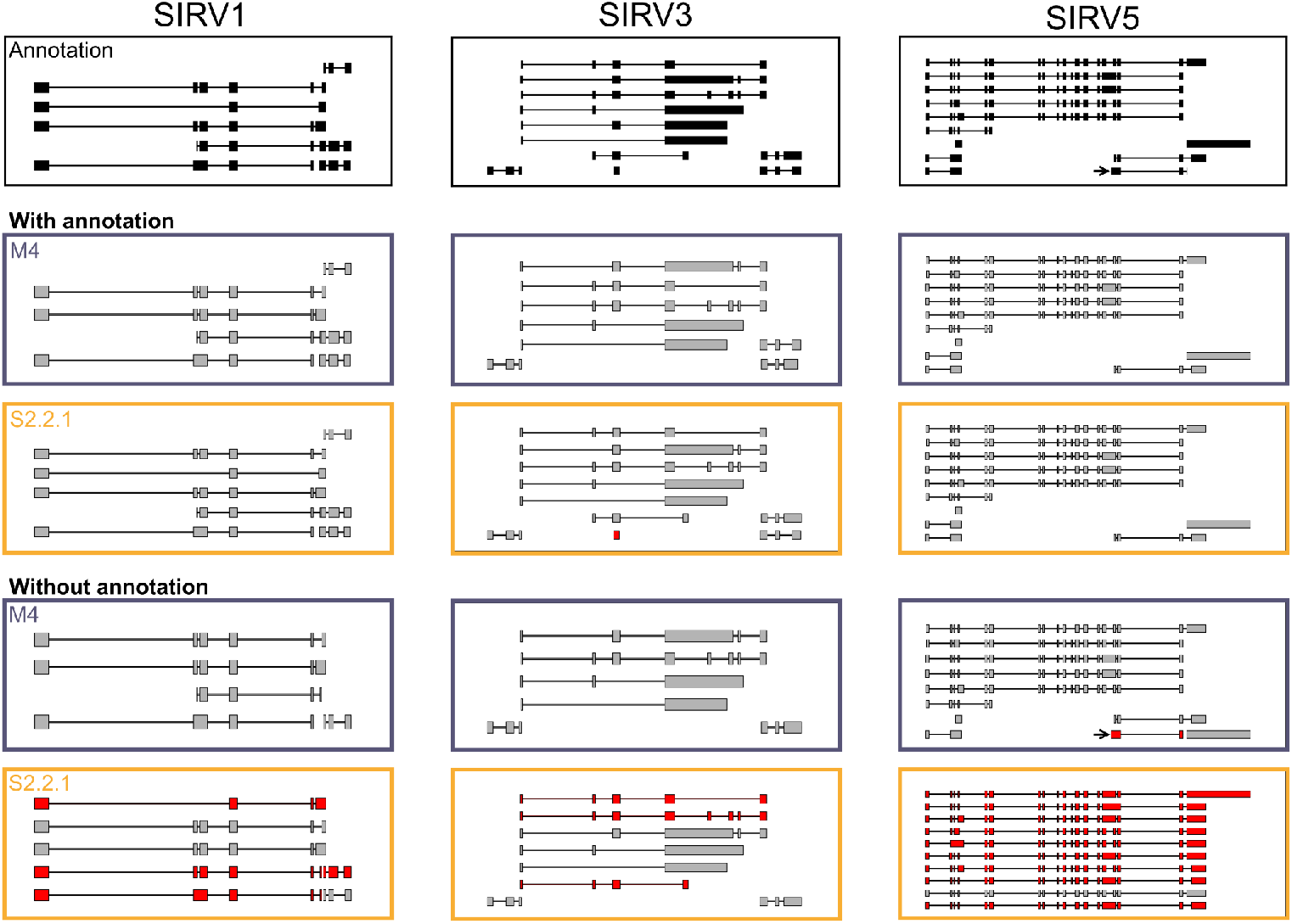
Mandalorion identifies SIRV isoforms with or without genome annotation. Genome browser views of isoforms identified by Mandalorion v4 (M4) and StringTie v2.2.1 (S2.2.1). True positive isoforms are shown in grey, False positive isoforms are shown in red. Mandalorion v4 performed well with and without annotation while StringTie v2.2.1 performance declined without annotation. Arrows highlight isoform SIRV503.

Overall, when no annotation file was available, Mandalorion continued to perform very strongly while StringTie performed worse in complex loci.

#### Evaluating Mandalorion v4 isoform quantification performance

In addition to identifying isoforms, Mandalorion also quantifies isoform abundance by counting the number of full-length sequencing reads associated with each isoform. To evaluate and compare the performance of Mandalorion v4 and StringTie v2.2.1 run with annotation, we focused our analysis on 23,884 and 21,156 isoforms correctly identified by Mandalorion v4 and StringTie v2.2.1 based on simulated PacBio data (SQANTI category FSM).

We compared the normalized read counts (transcripts per million - TPM) as determined by Mandalorion v4 and StringTie2 for these isoforms to the known TPM simulated for these isoforms. We found that Mandalorion v4 TPM values correlated very well with simulated TPM values, reaching a pearson r value of 0.992 compared to 0.942 reached by StringTie v2.2.1 (Fig. 4). We then performed the same comparison for the 21,814 and 20,539 isoforms identified by Mandalorion v4 and StringTie v2.2.1 without annotation. We found that Mandalorion v4 read TPM values continued to correlate very well with simulated TPM values reaching a pearson r value of 0.992 while the TPM correlation as determined by StringTie v2.2.1 actually improved to 0.986 (Fig. 4).

**Fig. 4:**
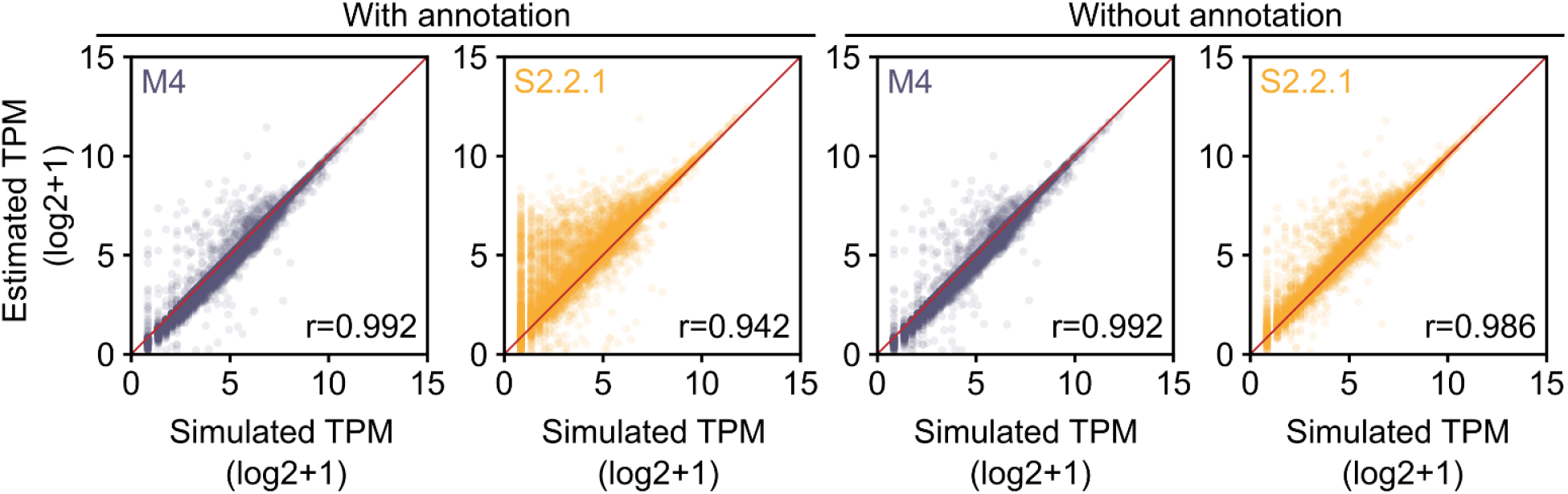
Mandalorion v4 quantifies isoform levels accurately in simulated data. Isoform levels (transcripts per million reads - TPM) for simulated isoforms as quantified by Mandalorion v4 (M4) and StringTie v2.2.1 (S2.2.1) are compared to the actual simulated TPM values as scatter plots. Isoform identification and quantification was performed with (left) and without (right) annotation.

In either case, Mandalorion v4 generated TPM values that correlated very closely with simulated TPM values.

#### Evaluating Mandalorion v4 isoform identification on UHR Iso-Seq data

Finally, we tested how Mandalorion v4 performed when analyzing publicly available Universal Human Reference (UHR) Iso-Seq data. This data set was released by PacBio and contains about 6.7 million full-length cDNA reads. Because there is no available ground truth for this data set, we focused on how the identified isoforms matched the GENCODE annotation and whether they reflected the actual read alignments they were based on.

With an annotation file available, Mandalorion v4 generated 53,348 isoforms of which 80.52% where categorized by SQANTI as either “full_splice-match” (FSM) or “novel_in_catalog” (NIC), i.e. it’s entire junction-chain (FSM) or all individual junctions (NIC) where present in the GENCODE v38 annotation (Fig. 5A). StringTie generated 93,611 isoforms of which 55.72% were categorized by SQANTI as either FSM or NIC. Using gffcompare[23], we compared Mandalorion v4 and StringTie v2.2.1 isoforms based on their junction chains (but not TSS and polyA sites). It is important to note that gffcompare removes isoforms with the same junction-chain within each sample before comparison and Mandalorion can generate multiple isoforms per junction-chain if there is read support for different TSS and polyA combinations for that junction-chain. After this removal step, 24,241 of the remaining isoforms were found to be shared between Mandalorion and StringTie. We then quantified how many reads mapped to each isoform shared between, or unique to each tool. For this, a read alignment had to perfectly match each splice site and its ends had to fall within a 30nt window (10nt upstream and 20nt downstream) of the isoform’s TSS and polyA site. Using this analysis we found that, in contrast to isoforms unique to Mandalorion, the majority of isoforms unique to StringTie had no read support (median of 0 reads mapped to the isoforms) (Fig. 5B). Even for the isoforms shared between Mandalorion and StringTie, Mandalorion isoforms had higher read support.

**Fig. 5:**
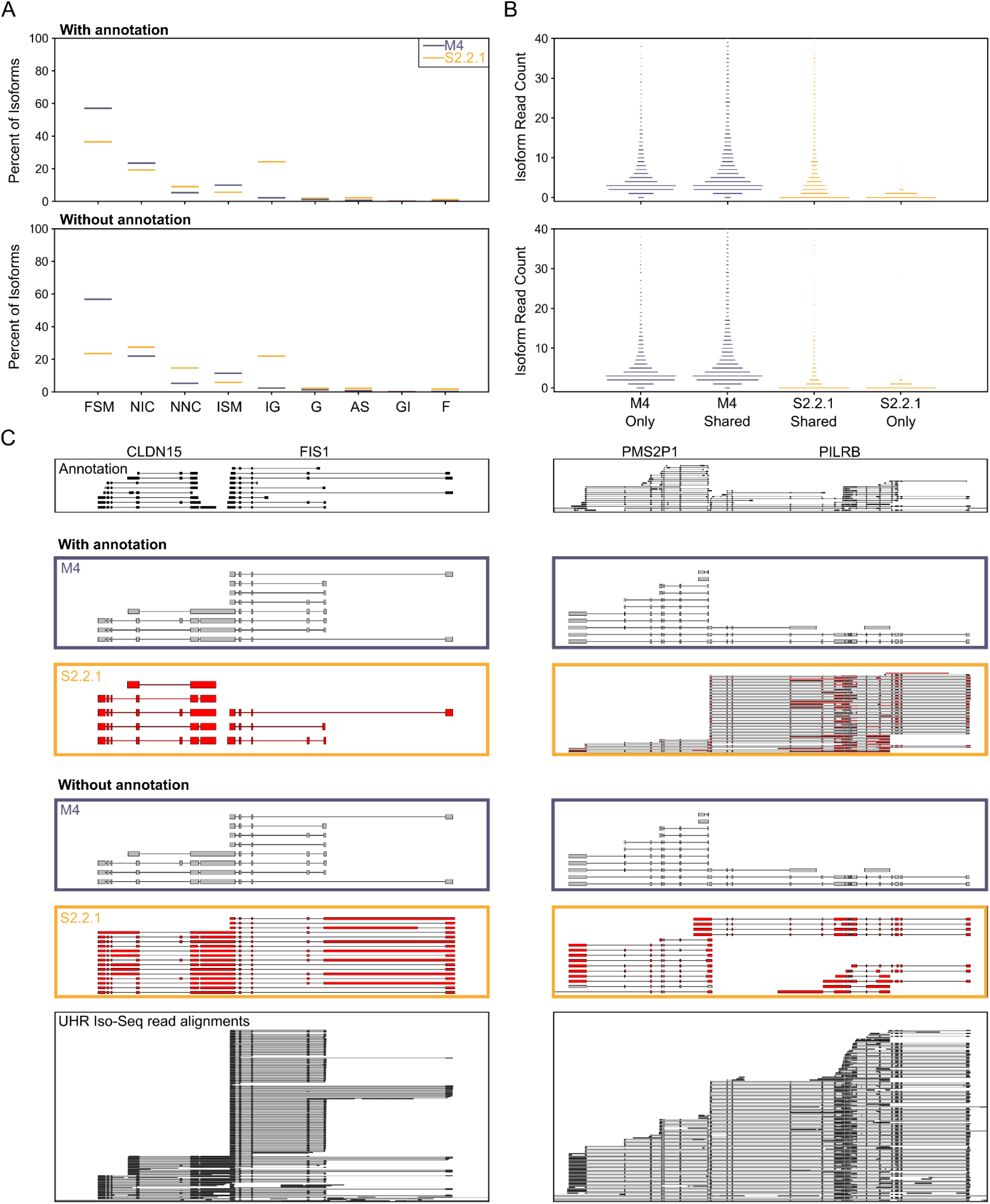
Mandalorion and StringTie performance on the Universal Human Reference Iso-Seq Dataset. A) SQANTI category composition of isoforms inferred by Mandalorion v4 (M4) and StringTie v2.2.1 (S2.2.1) with (top) and without (bottom). FSM=‘full-splice_match’, NIC=‘novel_in_catalog’, NNC=‘novel_not_in_catalog’, ISM=‘incomplete-splice_match’, IG=‘intergenic’, G=‘genic’, AS=‘antisense’, GI=‘genic_intron’, F=‘fusion’. B) Number of reads associated with isoforms exclusive to or shared between Mandalorion and StringTie. C) Genome Browser views of two genomic loci. Top to bottom, annotated isoforms, isoforms identified by Mandalorion v4 (M4) and StringTie v2.2.1 (S2.2.1) with or without annotation and PacBio Iso-Seq reads used for isoform identification are shown. Annotation and Iso-Seq read alignments are shown in black. Identified isoforms with or without read support are shown in grey or red.

Without an annotation file available, Mandalorion v4 generated 48,967 isoforms of which 78.74% were categorized as FSM and NIC. Using gffcompare we found that, after removing isoforms with the same junction-chain but different ends within each sample, 96.35% of the remaining 39,062 isoforms overlapped with the isoforms generated by Mandalorion run with annotation. StringTie generated 115,424 isoforms of which 51.00% were categorized as FSM and NIC. Interestingly, only 58.88% of these overlapped with the isoforms generated by StringTie run with annotation. We then, again, compared Mandalorion and StringTie isoforms and found that without annotation, even for isoforms that shared a junction-chain with a Mandalorion isoform, the median read support was 0 compared to 4 for the Mandalorion isoforms (Fig. 5B).

Overall this suggests two things. First, in contrast to StringTie, Mandalorion isoforms are very similar whether an annotation file is provided or not. Second, Mandalorion isoforms have TSS and polyA sites that more closely resemble the read alignments they are based on. We show two examples of this behavior (Fig. 5C) where StringTie isoforms with/without annotation are entirely different from each other and often not supported by any read alignments, while Mandalorion isoforms are highly similar with/without annotation and supported by read alignments.

Ultimately, this highlights that Mandalorion relies on the fact that the majority of Iso-Seq and R2C2 reads truly cover RNA transcripts end-to-end. We believe this is a strength when analyzing full-length cDNA data, but it also means that, in contrast to StringTie, Mandalorion cannot infer isoforms that may not be present in a full-length cDNA pool due to molecular biology or sequencing technology limitations, e.g. very long isoforms.

## DISCUSSION

We initially released Mandalorion in 2017 to identify isoforms based on the then fairly new full-length transcriptome data type [24]. Over the last 5 years we have continuously developed and used Mandalorion in several publications to analyze bulk and single cell data sets [9–11,25,26]. Version 4 is in many ways the culmination of our efforts over the last 5 years to turn Mandalorion from a hard to run collection of metaphorically duct-taped together scripts into an easy to install and run, robust, fast, and powerful tool. To highlight some improvements over previous versions: Mandalorion v4 now 1) requires only two non-standard python libraries (mappy, pyabpoa) and two standalone tools (minimap2, emtrey) to be installed, 2) can be run with concise input from the command line (see Methods), 3) is much faster (hours vs days) and requires less RAM due to optimized multithreading, 4) and has better Recall in poorly annotated loci.

Alongside Mandalorion, the full-length transcriptome sequencing field has matured as well and other tools have been designed for isoform identification based on this now established data type. These tools, which include but are not limited to FLAIR, IsoQuant, IsoTools, TALON, StringTie, Bambu, and FLAMES, present other approaches to the isoform identification problem and their “big-picture” differences can be compared in the LRGASP manuscript [19].

Here, we perform a separate, distinct analysis to show that Mandalorion represents a strong combination of Recall and Precision when analyzing ONT-based R2C2 or PacBio Iso-Seq data. In our comparison based on publicly available LRGASP and UHR data, Mandalorion compares favorably to the very powerful and highly used StringTie tool - especially in the absence of genome annotation. While running a tool entirely without genome annotation doesn’t reflect their likely usage on model organisms, it does allow us to predict performance in poorly annotated gene loci or in any transcriptome/genome combination that lacks a highly curated annotation, i.e. anything that isn’t a human or a mouse. Based on its performance here, we think Mandalorion is a powerful tool for de-novo genome annotation based on full-length transcriptome data.

One difference between Mandalorion and most other tools for the analysis of full-length transcriptome data is that Mandalorion generates a read-based consensus sequence for each isoform it identifies. This has already proven to be very useful for the analysis of HLA alleles [10] and should prove useful for future studies that involve different genotypes or haplotypes.

Overall, we believe that Mandalorion v4 is a strong addition to the toolbox of researchers analyzing full-length transcriptome data. As this data type becomes more common, additional tasks like variant detection and allele-specific isoform analysis will represent new challenges for tools like Mandalorion and represent exciting opportunities for further tool development.

## ACKNOWLEDGEMENTS

We want to thank Angela Brooks for reading and providing valuable feedback and advice with this manuscript. We acknowledge the efforts of all members of the LRGASP consortium with special thanks to everybody involved in generating, simulating, organizing, and hosting data. We acknowledge funding by the National Institute of General Medical Sciences / National Institutes of Health Grant R35GM133569 (to C. V.).

## DATA AVAILABILITY

Part of the data used in this manuscript was generated for the LRGASP consortium by several labs, including ours (for R2C2 data). All this data is publicly available. R2C2 and PacBio data for the Mouse F121-9 cell line can be found at ENCODE under file IDs ENCFF513AEK, ENCFF824JVI, ENCFF104DMI, ENCFF850MIB, ENCFF412UHU, ENCFF595TIH for R2C2 data and ENCFF874VSI, ENCFF667VXS, ENCFF313VYZ for PacBio data. Simulated mouse data is available as part of the data provided by the LRGASP consortium at Synapse under ID syn25683381. PacBio Iso-Seq UHR data is available at https://downloads.pacbcloud.com/public/dataset/UHR_IsoSeq/

## CODE AVAILABILITY

Mandalorion v4 is available at https://github.com/christopher-vollmers/Mandalorion.

## METHODS

### Read preprocessing

#### R2C2

~~~
python3 Mandalorion/utils/remove_polyA.py-i input.fasta -o output.trimmed.fasta -t 68,50
~~~

#### Simulated PacBio

~~~
python3 Mandalorion/utils/removePolyA_nonDirectionalInput.py -i input.fasta -o output.trimmed.fasta -t 1,1
~~~

### Mandalorion

#### With annotation

~~~
python3 Mando.py -p ./ -f reads.fofn -W basic,SIRV -G lrgasp_grcm39_sirvs.fasta -t 50 -I 200 -g lrgasp_gencode_vM27_sirvs.gtf
~~~

#### Without annotation

~~~
python3 Mando.py -p ./ -f reads.fofn -G lrgasp_grcm39_sirvs.fasta -t 50 -I 200
~~~

### StringTie

For StringTie we used the alignments generated by running Mandalorion after sorting and converting to bam using samtools[27]

#### With annotation

~~~
*stringtie mm2Alignments*.*sorted*.*bam -o stringtie_annot*.*gtf -L -p 50 -G lrgasp_gencode_vM27_sirvs*.*gtf*
~~~

#### Without annotation

~~~
*stringtie mm2Alignments*.*sorted*.*bam -o stringtie_annot*.*gtf -L -p 50*
~~~

### SQANTI analysis

#### R2C2, PacBio, and Freestyle LRGASP data

~~~
python3 sqanti3_lrgasp.challenge1.py isoform.gtf lrgasp_gencode_vM27_sirvs.gtf lrgasp_grcm39_sirvs.fasta --json experiment.json
--cage_peak refTSS.mouse.bed --polyA_motif_list polyA_list.txt -c ES_Illumina_STARpass1_SJ.out.tab -d ./SQANTI -o output --gtf
~~~

#### Simulated PacBio LRGASP data

~~~
python3 sqanti3_lrgasp.challenge1.py isoform.gtf simulated_isoforms.gtf lrgasp_grcm39_sirvs.fasta --json experiment.json --cage_peak
refTSS.mouse.bed --polyA_motif_list polyA_list.txt -c ES_Illumina_STARpass1_SJ.out.tab -d ./SQANTI -o output --gtf
~~~

#### PacBio UHR data

~~~
python3 sqanti3_lrgasp.challenge1.py isoform.gtf lrgasp_gencode_v38_sirvs.gtf lrgasp_grch38_sirvs.fasta --json experiment.json
--cage_peak refTSS.human.bed --polyA_motif_list polyA_list.txt -c WTC11_Illumina_STARpass1_SJ.out.tab -d ./SQANTI -o output --gtf
~~~

Samtools [27], Numpy [28,29], Scipy [30], and Matplotlib [31] libraries were used extensively for analysis.

---

**Table S1.:** Recall and precision of Mandalorion and StringTie versions.

| Tool | Ground Truth | Read source | Reference annotation | Recall | Precision |
| --- | --- | --- | --- | --- | --- |
| Mandalorion v4 | Simulated isoforms | IsoSeqSim | Yes | 87.96% | 92.23% |
| Mandalorion v3.6 | Simulated isoforms | IsoSeqSim | Yes | 85.12% | 91.80% |
| StringTie v2.2.1 | Simulated isoforms | IsoSeqSim | Yes | 77.91% | 91.86% |
| Mandalorion v4 | Simulated isoforms | IsoSeqSim | No | 80.34% | 91.59% |
| Mandalorion v3.6 | Simulated isoforms | IsoSeqSim | No | 66.95% | 89.50% |
| StringTie v2.2.1 | Simulated isoforms | IsoSeqSim | No | 75.64% | 90.86% |
| Mandalorion v4 | SIRV isoforms | R2C2 | Yes | 88.40% | 98.39% |
| StringTie v2.2.1 | SIRV isoforms | R2C2 | Yes | 84.06% | 95.08% |
| Mandalorion v4 | SIRV isoforms | R2C2 | No | 76.81% | 98.15% |
| StringTie v2.2.1 | SIRV isoforms | R2C2 | No | 42.03% | 55.77% |
| Mandalorion v4 | SIRV isoforms | PacBio | Yes | 89.86% | 100% |
| StringTie v2.2.1 | SIRV isoforms | PacBio | Yes | 91.30% | 96.92% |
| Mandalorion v4 | SIRV isoforms | PacBio | No | 71.01% | 98% |
| StringTie v2.2.1 | SIRV isoforms | PacBio | No | 36.23% | 53.19% |
| Mandalorion v4 | SIRV isoforms | Combined | Yes | 94.20% | 98.48% |
| StringTie v2.2.1 | SIRV isoforms | Combined | Yes | 84.06% | 95.08% |
| Mandalorion v4 | SIRV isoforms | Combined | No | 79.71% | 96.49% |
| StringTie v2.2.1 | SIRV isoforms | Combined | No | 42.03% | 50% |

